# Electrophysiological “Brain Power” Dissipation: MEG Lower-Bound Estimates and Regional Metabolic Correlates

**DOI:** 10.1101/2025.09.29.679241

**Authors:** Vikram Nathan, Malte Hoeltershinken, Eleanor Hill, Carsten Wolters, Sylvain Baillet

## Abstract

The human brain is often described as remarkably energy efficient, with its metabolic power consumption estimated at approximately 20 watts (W). Biophysical models partition this metabolic budget across cellular processes, including neuronal signaling. However, the electrical power dissipated by the synchronized cortical activity measurable with noninvasive electrophysiology remains unknown and should not be equated with total cortical energetic demand.

Here, we used magnetoencephalography (MEG) source imaging of resting-state activity to estimate electrical power dissipation in cortical gray matter. Across 30 participants, mean total primary power summed over the modeled cortical source space was 6.6 × 10□□ W. We interpret this value as an empirical lower-bound estimate for the synchronized component of cortical activity accessible to MEG, rather than as an estimate of total cortical power. Targeted sensitivity analyses and order-of-magnitude checks suggested that the tested methodological choices could shift the estimate by approximately one to two orders of magnitude, but not enough to bridge the gap with metabolic benchmarks. Under a magnitude-based root-mean-squares approximation, finite-element estimates of secondary-current dissipation were approximately 30% of primary power in four central-tendency exemplars.

Spatial patterns of MEG-derived power showed associations with PET-derived oxygen metabolism and transcriptomic maps of ketone metabolism, with heterogeneous effects across functional networks. These results define the scope and limitations of MEG-derived electrophysiological power estimates and motivate multiscale models linking synchronized electrical activity to regional metabolism.

**Key Points:**

1. Across 30 participants, mean total primary power summed over the modeled cortical source space was 6.6 × 10□□ W.
2. Phantom, inverse-solution, resistance, and FEM analyses provide order-of-magnitude sensitivity checks rather than direct validation of absolute power.
3. Regional power maps showed spatial associations with PET-derived oxygen metabolism and transcriptomic indices of ketone metabolism after spatial-null correction.

## Introduction

The human brain processes information with remarkable power efficiency (approximately 20 W), requiring less power than a typical laptop computer (Balasubramanian, 2021). This estimate of total power demand is primarily derived from stoichiometric calculations of oxygen and glucose consumption measured by positron emission tomography (PET) and adenosine triphosphate (ATP) turnover determined by ³¹P magnetic resonance spectroscopy imaging (Clarke & Sokoloff, 1999; Zhu et al., 2012). Despite this general understanding, precisely quantifying the energetic costs of distinct neuronal processes remains challenging.

Mathematical “power budget” models based on electrophysiological properties of neurons have offered theoretical partitions of this consumption. Such models translate measured ionic fluxes into power expenditures associated with distinct neural processes, for example, differentiating between white versus gray matter signaling, and action potential conduction versus passive membrane properties (Attwell & Laughlin, 2001; Lennie, 2003). Levy & Calvert (2021) further partitioned gray matter activity into “communication” and “computation,” showing computation to be ∼35 times more efficient and contributing ∼0.1 W of the brain’s overall power. However, it remains unclear to what extent such theoretical power budgets can be related to the synchronized component of human cortical activity measurable with whole-cortex electrophysiology.

Previous work indicates that postsynaptic potentials constitute a substantial portion of gray matter’s energetic demand, largely due to sodium and potassium currents associated with synaptic transmission (Alle et al., 2009). At the whole-cortex scale, electrophysiological and hemodynamic recordings in non-human primates have demonstrated that local field potentials (LFPs), primarily reflecting postsynaptic potentials, predict resting-state hemodynamic fluctuations better than action potentials (Logothetis, 2008). Yet, the electrical power dissipated by large-scale synchronized postsynaptic activity detectable with noninvasive electrophysiology remains unknown.

To address this gap, we developed a data-driven approach leveraging the temporal resolution and cortical sensitivity of magnetoencephalography (MEG; Baillet, 2017). Specifically, we estimated electrical power dissipation associated with MEG-visible, synchronized cortical current fluctuations during rest. By mapping primary current dipoles to vertex-associated cortical patches, we derived a regional power metric that we interpret as a lower bound on the synchronized activity accessible to MEG, not as an estimate of total cortical power.

## Methods

### MEG and MRI Data Acquisition

Resting-state MEG data were available from 31 healthy adults (age range: 21–47 years; 14 women), recorded during eyes-open rest with a CTF 275-channel system at 2,400 Hz. Although more than five minutes of data were available for each participant, all analyses used the first 120 seconds of each recording (see Temporal Aggregation of MEG Primary Current Dipoles). Data were recorded with an anti-aliasing low-pass filter below 600 Hz. Individual head shape, captured at approximately 100 scalp locations, and fiducial points (nasion, left and right preauricular points) were digitized with a Polhemus Fastrak system using Brainstorm (Tadel et al., 2011) and co-registered to T1-weighted MRI volumes processed in FreeSurfer 5.3 for tissue segmentation and cortical surface mesh triangulation (Fischl, 2012).

### FEM Model Creation and Forward Modelling

To estimate currents on the cortical surface (which we refer to as “sources”) from MEG sensors, we used a standard two-step approach. Our first step was forward modelling, which estimates the “lead-field” matrix that predicts MEG sensor activity from cortical sources. Our second step is inverse modelling, which uses mathematical inference and the lead-field matrix to estimate the cortical sources that generate a given MEG recording and fulfill the prior assumptions of the chosen inverse approach (Baillet, Mosher, & Leahy, 2001).

Recent work has shown that forward modelling is enhanced by detailed estimates of the anatomy underlying cortical sources, producing a “volume conductor” that captures spatial complexities when relating source to sensor activity (Schrader et al., 2021). Although classical forward modelling approaches, such as assuming an overlapping-spheres geometry, produce faster closed-form solutions, we used the finite element method (FEM) forward model to remain methodologically consistent between primary and secondary power estimations (see Cortical Patch vs. FEM-Derived Power) and to support spatial comparisons with whole-cortex metabolic maps.

Surfaces for the head, outer skull, and inner skull were generated from each participant’s T1 MRI using *Brainstorm*’s boundary element model implementation. *Iso2Mesh* (Tran et al., 2020) was then used to generate the FEM model, with the “*MergeMesh*” option selected to tessellate internal cavities. Default conductivity values were set to 0.33 S/m for brain, 0.43 S/m for scalp, and 0.008 S/m for skull. For all n=31 participants, this procedure successfully produced three-layer models (∼51,700 vertices and ∼276,000 elements; Figure 3C). FEM modeling enabled estimation of secondary currents across tissue layers, complementing primary dipole-based estimates.

We then used these FEM models for FEM forward modelling in *DUNEuro* to produce a lead-field matrix for each participant (Medani et al., 2023; Schrader et al., 2021). We used the default settings within *Brainstorm* for the FEM solver. The FEM model itself was generated with *Iso2mesh* (Fang & Boas, 2009), using the default maximum tetrahedral element volume (0.1 cm³) and 100% element-keep ratio after mesh simplification. For the *DUNEuro* solver configuration, we used a fitted, isotropic FEM formulation with a conjugate-gradient (CG) solver and the Venant source model. The cortical source space was not shrunk inward from the pial/central surface before lead-field computation and was not artificially forced into gray matter, and no anisotropic/DTI-derived conductivity tensors were used.

The FEM forward model maps cortical dipole moments, expressed in A·m, to sensor-level magnetic fields, expressed in T. The lead-field matrix G therefore has units of T/(A·m). Each participant had N□ = 15,002–15,011 source locations, with three orientation components per location; consequently, G had dimensions Nc × 3N□, where Nc = 268–274 MEG channels varied slightly by participant (modal value 270), or approximately 270 × 45,000. The corresponding inverse operator maps sensor fields to dipole moments and has units of (A·m)/T.

### MEG Source Mapping

Preprocessing followed good-practice guidelines (Gross et al, 2013), including artifact (e.g., cardiac and ocular) removal by signal-space projection (SSP) available in *Brainstorm*, ensuring the reliability of our MEG sensor data for source estimation. Environmental noise covariance was estimated from two-minute empty-room recordings acquired immediately before each MEG session.

Source reconstruction was performed using minimum norm estimation (MNE) implemented in *Brainstorm*. MNE is a mathematical technique that multiplies the inverse operator M by the MEG sensor data (in T for each source) to obtain primary current dipoles (which we refer to as I_dip_ in units of A.m.) for each of the ∼15,002 cortical sources. The inverse operator M implemented in Brainstorm is defined below (Hämäläinen 2005):

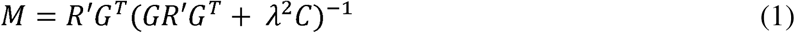

where R′ is the modeled source covariance matrix, G is the lead-field matrix, C is the noise covariance matrix, and *λ*² is the regularization parameter. Brainstorm parameterizes MNE regularization through an assumed signal-to-noise ratio (SNR), with *λ*² = 1/SNR². The primary analyses used the Brainstorm default of SNR = 3 (*λ*² = 1/9); sensitivity analyses used SNR = 1 and SNR = 5 (*λ*² = 1 and 1/25, respectively).

Smaller *λ*² values impose less shrinkage and generally yield larger reconstructed dipole moments, whereas larger values impose stronger regularization. Because reconstructed amplitude directly affects power, we recomputed both current-dipole and vertex-wise power estimates under each SNR setting.

We chose to use MNE over other source estimation techniques due to several key advantages. Firstly, rather than estimating a relative, statistical map of currents across the cortex, which is the case when using other methods like dSPM or sLORETA, MNE offers physical units for the estimated current dipoles, which was necessary for our biophysical modelling. Secondly, the MNE solution can resolve complex spatial sources with similar spatial precision to dSPM and sLORETA (Hauk et al., 2011). Finally, particularly in the context of resting-state data, MNE outperforms dipole fitting and scanning methods, which typically result in a large amount of unexplained variance due to difficulties in choosing the number of sources (Gramfort et al., 2014).

We used MNE with default hyperparameter settings in Brainstorm (Tadel et al., 2011), noting that regularization may smooth focal sources over a wider area. Source locations were restricted to the central cortical surface, but dipole orientation was unconstrained, yielding three moment components at each vertex. We used unconstrained orientation to avoid imposing a strictly surface-normal direction in the presence of local geometric and segmentation uncertainty. Because this choice affects reconstructed amplitude, we repeated the primary analysis with dipoles constrained normal to the cortical surface.

Because the conversion I = D/L uses cortical thickness as an effective current-path length, it is most directly interpretable for the surface-normal dipole component. For free-orientation solutions, D/L should therefore be understood as an effective patch-level current scale rather than a literal translaminar current. We assessed sensitivity to this approximation using surface-normal constrained sources.

### Current-Dipole Recovery with Physical Phantoms

To characterize current recovery, MEG data were analyzed from two calibration phantoms: the CTF phantom (a single dipole in a conducting sphere) and the Elekta phantom (32 open-air dipoles arranged beneath a hemispherical surface; Figure 2A)

The CTF phantom generated dipole moments of 1,800 and 180 nA·m at 7 and 23 Hz, respectively, sampled at 2,400 Hz. For the Elekta analysis, of the amplitudes available, we only used the tutorial’s 200 nA·m recording to compare against the CTF phantom’s lower-amplitude condition, comprising sequential recordings from 32 dipoles oscillating at 20 Hz, acquired at 1,000 Hz and resampled to 100 Hz. The tutorial also distributes 20 and 2,000 nA·m recordings; these conditions were not analyzed, and their earlier inclusion in the Methods was a transcription error rather than evidence of omitted analyses. Source datasets are available through the Brainstorm CTF and Elekta phantom tutorials (https://neuroimage.usc.edu/brainstorm/Tutorials/PhantomCtf; https://neuroimage.usc.edu/brainstorm/Tutorials/PhantomElekta).

For the source mapping of these phantom currents, we used a similar MNE approach, with the exception of the overlapping spheres forward model, due to the lack of underlying anatomy for the phantom data. First, we segmented data into epochs surrounding the activation of each emitted dipole. The epochs were 140 ms in length for the CTF phantom recordings, and 400 ms for the Elekta phantom, respectively; these epoch lengths were chosen based on existing *Brainstorm* tutorials (see above web links), and were meant to capture one or more full oscillatory cycles. Next, for each phantom dipole, we averaged the MEG data across all epochs to reduce noise before our MNE source reconstruction.

For each phantom dipole’s epoch-averaged activity, we then identified the time points of peak current emission (note: there is one dipole for CTF, and 32 separate dipoles for Elekta). We then calculated and summed up the MNE-reconstructed amplitudes across the volume of each phantom at each of these peak time points, arriving at the total current strength reconstructed for each emitted dipole.

This epoching and averaging procedure yielded three “peak” current estimates for the 1800 nA·m current dipole and seven peaks for the 180 nA·m dipole from the CTF phantom. There was only one peak (60 ms) for the Elekta phantom, but the source could be localized separately at this peak for each of the 32 dipoles.

The phantom analyses characterize current-dipole recovery rather than overall power dissipation because the phantom sources lack resistances comparable to cortical tissue. The CTF phantom is immersed in saline of unknown conductivity, whereas the Elekta sources are surrounded by air; MRI-derived cortical thickness and support area are also undefined in these configurations. We therefore used the phantom data as order-of-magnitude checks of current-amplitude recovery, which directly affects inferred power, rather than as validation of absolute cortical power.

### Cortical Patch vs. FEM-Derived Power

#### (i) Primary Cortical Patch Power

Each cortical source vertex was associated with a local surface support region, which we term a vertex-associated cortical patch. These patches are numerical support regions of the source mesh and should not be interpreted as canonical biological cortical columns (Molnár et al., 2020). We estimated power dissipation separately for each cortical patch using Ohm’s law, with current amplitudes derived from MEG source reconstructions and resistance from co-registered structural MRI data. Neural impedance was assumed to be purely resistive, neglecting capacitive and inductive components (Baillet et al., 2001).

The downsampled version of the Freesurfer surface mesh (15,002 vertices) was used to define vertices on the cortex, as well as the support area (A) attached to each vertex. CAT12 was used to calculate cortical thickness (L) at each vertex due to the increased efficiency of its implementation in *Brainstorm*.

Resistance (*R*) was then calculated as:

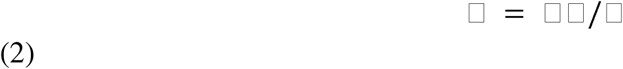

where ⍰ is cortical resistivity (3.5 Ω.m; Abboud et al., 2021); *L* is cortical thickness (taken as dipole length, in meters), and *A* is surface area, defined as one-third of the area of all the neighboring mesh triangles (Hillebrand & Barnes, 2002), in square meters.

Power dissipation (*P*), in Watts (W), at each cortical vertex was then defined as:

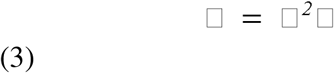

Let D□, in A·m, denote the reconstructed dipole-moment vector at vertex v. Approximating the effective patch current as I□=□D□□/L□, the power dissipated at that vertex is

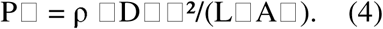

Total primary P was computed by calculating P□ separately at every vertex and then summing across all modeled cortical patches.

##### (ii) Secondary Current Power (FEM)

Secondary volume currents in head tissues (brain, skull, scalp) were estimated using FEM-derived electric fields (*E*). After computing the electric field *E*, in V/m, due to neural activity, the associated secondary currents’ density (*j^S^*) in A/m^2^ was defined as:

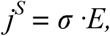

where σ denotes tissue conductivity in S/m (Hämäläinen et al., 1993). The corresponding power dissipation was then computed as the integral of the electric field times the current density vectors across all volumetric sources:

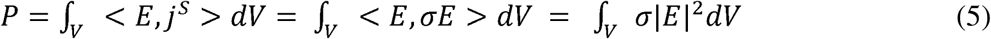

where ⟨E,j⟩ denotes the inner product (Griffiths, 2023, Chapter 8.1.2).

FEM simulations were performed using the Python interface of the *DUNEuro* toolbox (Schrader et al, 2021; Höltershinken et al., 2025). A continuous Galerkin approach with piecewise affine trial and test functions was applied, with partial integration used to handle dipole singularities.

For each source i, the FEM input moment was the component-wise RMS vector

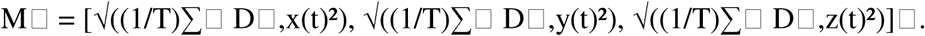

This construction preserves the time-averaged squared magnitude of the primary dipole but removes the sign and temporal covariance of its orientation components. Consequently, it does not represent an instantaneous physical dipole orientation. Moreover, summing the fields generated by these RMS vectors before computing total secondary P may introduce artificial cross-source coherence. We therefore interpret the secondary-power results as a magnitude-based approximation and sensitivity analysis, rather than an unbiased estimate of time-averaged secondary dissipation.

For each dipole at position x□ with moment M□, the FEM solver generated a corresponding field vector E□. By linearity of the governing equations, the total field was:

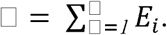

We then computed two metrics:

Total secondary power, which summates the secondary currents from all primary sources:

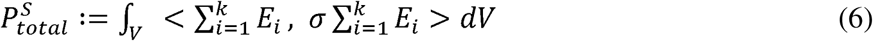

Secondary dipole power, which summates P from each primary dipole’s secondary currents:

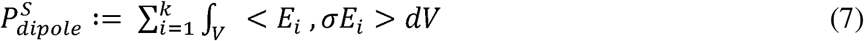

Finally, note that in the special case of a current *I* passing through a resistor *R*, the general FEM formulation reduces to the Ohmic definition in Equation 3. Thus, this framework generalizes the primary cortical patch model to more complex, anisotropic conductors.

### Temporal Aggregation of MEG Primary Current Dipoles

Before calculating the power metrics, we summarized each participant’s source activity as a component-wise root-mean-square (RMS) current-dipole vector over the first 120 seconds of the recording, using the following formula:

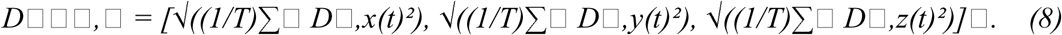

Here, the vector D□□□,□ is the component-wise RMS current dipole at each vertex v, T is the total number of recorded time points (2,400 Hz × 120 s), and D denotes the x, y, or z component of the source-estimated primary dipole.

This duration produces a stable estimate of the average current amplitude (Wiesman et al., 2022), and the resulting RMS current dipoles can be used as summary metrics for calculating the primary and secondary P. We note that substituting D_rms, v_ into Equation (4) is equivalent to calculating the average primary dipole P across all T timepoints’ *D_i_* values, since all other quantities in Eq. 5 are time-invariant (see Supplementary Methods 1).

Although equivalent for primary dipole P, we note that using *D_rms, v_* is an approximation for estimating the average secondary dipole P and total secondary P over time. To estimate secondary currents, the DUNEuro FEM solver uses linear operations to produce a vector of electrical fields *E_i_* from primary current dipoles *i*; hence, the temporal average of the estimated electrical fields is not the same as estimating a single set of electrical fields due to the RMS primary current dipole, since the RMS applies a nonlinear transformation to the currents.

### Dendritic vs. Gray Matter-Derived Resistances

Our “primary dipole P” framework estimates electrical resistance within each cortical patch by assuming uniform gray-matter resistivity, scaled by cortical thickness and surface area. However, because cortical tissue also includes non-neuronal components, we derived a complementary estimate of resistance attributable specifically to ensembles of parallel apical dendrites within a given cortical patch. This analysis served to examine whether the order of magnitude of our resistance estimates was changed by accounting for the biophysical properties and neuron counts in each patch.

To approximate the resistance of Layer 5 pyramidal neurons (the cell population most often considered the principal generator of MEG signals) (Murakami and Okada, 2006), we used established values from Kasevich and LaBerge, 2011 for dendritic resistivity (29.7 ohm-cm), length (1200 µm), and cross-sectional area, assuming a circular cross-section with a 5 µm diameter. Single-neuron resistance was calculated from Equation 2, yielding R_neuron_=1.816*10^7^ Ω.

We then estimated the number of apical dendrites contributing to the signal of each cortical patch using the following procedure (see Supplementary Figure 1 for a schematic):

1. <u>Parcel assignment:</u> Each cortical vertex was assigned to a Desikan-Killiany-Tourville (DKT) atlas parcel based on *FreeSurfer 5.3* parcellations (Fischl, 2012).
2. <u>Neuronal proportion:</u> The normalized neuronal cell-type proportion (*p_neur_*) for each parcel was extracted from whole-brain maps derived by deconvolving Allen Human Brain Atlas bulk-expression data (Pak et al., 2024). In the source study, each voxel’s cell-type proportions were estimated using genetic markers for six cell types, and scaled relative to gray-matter density within each voxel; *p_neur_* is the neuronal volume fraction from this estimate.
3. <u>Density scaling:</u> *p_neur_* was multiplied by published neuron density estimates (neurons/mm^2^) for gross anatomical regions (e.g., frontal, temporal, occipital; see Klein and Tourville (2012)), derived from DAP staining (Ribeiro et al., 2013).
4. Patch scaling: This density was multiplied by the support area (*A*) in mm^2^ at each vertex to yield the total number of neurons (*N*) per cortical patch.

Assuming apical dendrites behave as resistors in parallel, the total effective resistance in a cortical patch was given by:

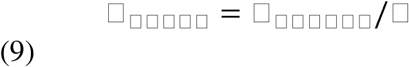

We applied both resistance formulations to all 30 participants in the primary-analysis cohort. Because both formulations contain the support-area factor 1/A, we also assessed their spatial correspondence after algebraically removing this shared term, thereby comparing ρL with the corresponding neuron-density-dependent component of the dendritic formulation.

### Six-Layer FEM Model

To compare the relative proportion of currents within our three-layer FEM model against a more anatomically precise six-layer FEM model, we simulated a 20 nA.m somatosensory evoked field (SEF) in the grey matter compartment of a publicly available six-layer FEM model for a different representative subject (n=49 years old, male). Briefly, this individual had three images; T1w- and T2w MRI, and diffusion tensor imaging (DTI), which were used to segment the head into six compartments. These compartments were then used to delineate the elements of a six-layer, volumetric, tetrahedral model with 885,214 nodes and 5,335,615 tetrahedrons, using the CGAL software (The CGAL Project, 2013) within iso2mesh. For full information into the FEM model creation methods, see https://zenodo.org/records/3888381 (Piastra et al., 2021); note that we did not use the recordings from this repository, and instead simulated the 20 nA.m SEF.

### Metabolism Correlations

We sought to compare our new metric of electrophysiological power dissipation against various spatial maps of cortical metabolism. We used the *neuromaps* toolbox (Markello et al., 2022) to access publicly available PET datasets (Vaishnavi et al., 2010) of glucose metabolism, measured using [^18^F]-labeled fluorodeoxyglucose (FDG) PET, and oxygen metabolism, measured with three scans involving the measurement of [^15^0]-labeled water, carbon monoxide, and oxygen. Similarly, we compared MEG-derived maps to gene-derived maps of metabolic processes, derived from whole-cortex microarray gene expression in six postmortem subjects from the Allen Human Brain Atlas (Hawrylycz et al., 2012). For each of 34 metabolic processes, spatial maps were generated by averaging the expression of genes within the corresponding Gene Ontology term (Pourmajidian et al., 2025). Prior to calculating correlations, all of the MEG, PET, and gene-derived data were parcellated into the Schaefer-600 atlas (Schaefer et al., 2018).

We derived p-values using a spatial permutation (’spin’) test with 10,000 rotations (Alexander-Bloch et al., 2018) to derive a null-distribution that was sufficiently granular to assess our Bonferroni-corrected significance thresholds. Parcel centroids of one spatial map were projected onto a sphere and randomly rotated; each parcel was reassigned the value of whichever parcel fell nearest to it under that rotation, generating a spatially-permuted null map that preserves spatial autocorrelation. This null map was correlated against the second, unrotated map to build a null distribution, and p-values were calculated as the proportion of null correlations at least as extreme as the observed correlation.

Since we ran multiple comparisons across types of metabolism and networks, we applied Bonferroni corrections within each family of comparisons. We note that there are different Bonferroni corrections to the critical value □=0.05 for whole-brain PET correlations (n=2 comparisons), gene-derived metabolism correlations (n=34 comparisons), and within-network correlations (n=7 comparisons), separately applied to PET and gene-derived measures. Additionally, although we used the same Bonferroni-corrected critical value for all comparisons within a family, each spatial map had a distinct null distribution, resulting in a different critical Spearman correlation for each spatial map.

Finally, beyond our comparisons with current-dipole correlations, we ran a partial correlation analysis to establish whether our correlations between P and metabolic maps are driven by the underlying input features. Using the same spatial null modelling and Bonferroni correction as outlined earlier, we separately calculated the partial correlation between P and each metabolism map, but assessing cortical thickness (L), support area (A), and their combination within primary dipole P (proportional to 1/(L×A)) as separate covariates.

## Results

### Initial MEG-Derived Cortical Power Estimate

We processed 31 recordings. Participant sub-0011 was excluded from the primary analysis because total primary P was 16.4 standard deviations above the mean of the remaining participants (1.16 × 10□□ W). The sensitivity analysis retaining this participant is reported in Supplementary Figure 4. All primary summary statistics therefore refer to the remaining n=30 participants.

Across the n=30 primary-analysis participants, mean total primary P summed over the modeled cortical source space was 6.6 × 10□□ W, with individual values ranging from 2.7 × 10□□ to 3.8 × 10□□ W. These values reflect the combination of cortical currents, thickness, and resistivity used in our modeling (Figure 1B).

**Figure 1:**
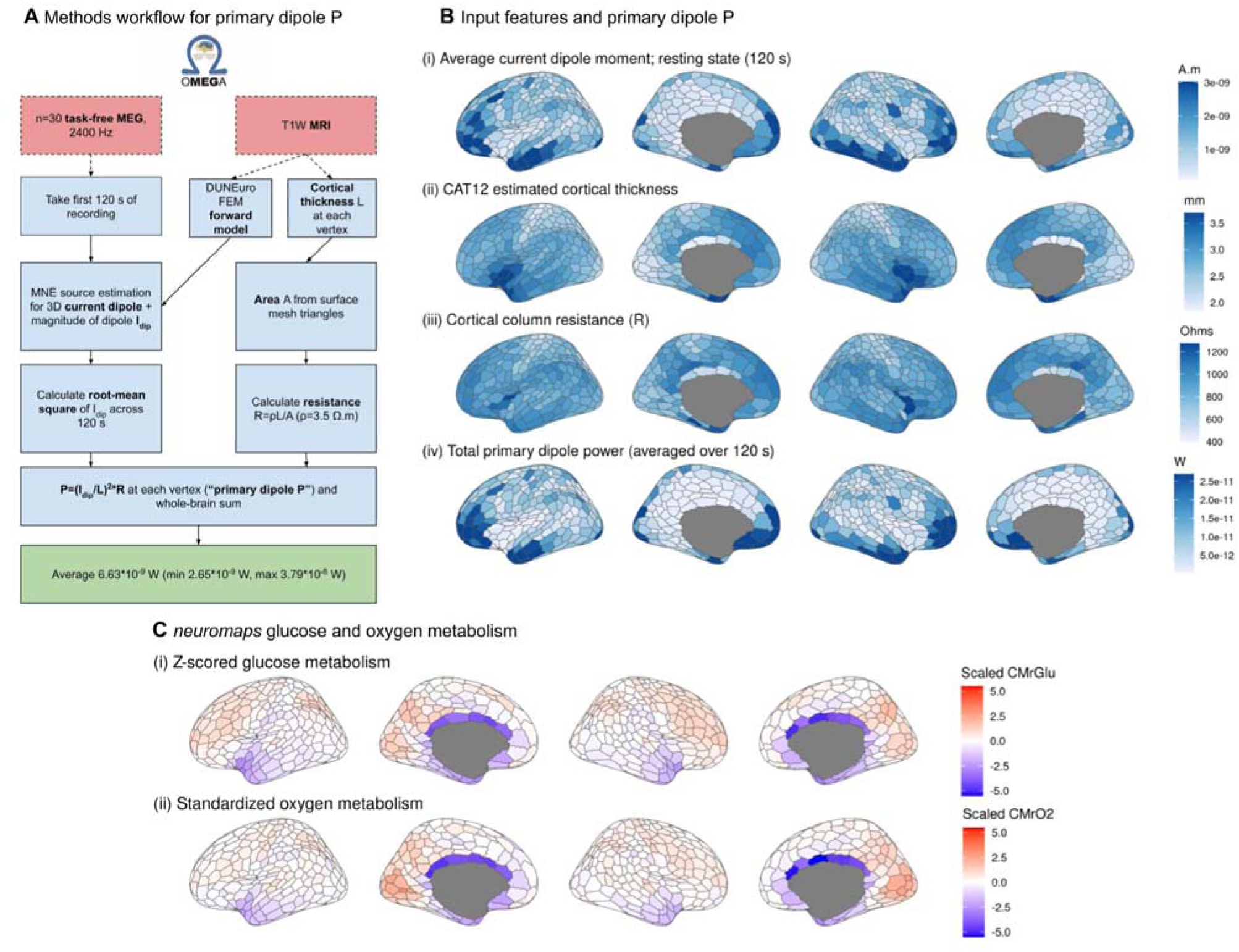
Methods workflow (Fig. 1A) and input features (Fig. 1B) used to calculate power at primary current dipoles (primary P). In Fig. 1A, ** indicates that sub-0011 was excluded from primary summary statistics as described in the Results. Visualizations in Fig. 1B–C use the Schaefer-600 parcellation. Figure 1B shows the MNE current dipole, CAT12 cortical thickness, cortical-patch resistance, and resulting mean primary-P map across the n=30 primary-analysis participants. Figure 1C shows glucose- and oxygen-metabolism maps from n=33 individuals (Vaishnavi et al., 2010) accessed through neuromaps. Outliers beyond 1.5 times the interquartile range were aggregated and all values were z-scored for visualization only; neither operation was used for the inferential correlations. The parcel “7Networks_Default_pCunPCC_14” was unavailable because of a Brainstorm parcellation issue.

### Order-Of-Magnitude Checks for Current Dipole Estimates and Resistive Modeling

To determine whether underestimated source currents might explain the low power values, we evaluated the performance of MNE-based source reconstruction using MEG recordings from physical phantoms with known current dipoles, testing three conditions across two phantom systems. For each phantom condition, we calculate the sum of Euclidean dipole norms across all estimated sources, which could yield a recovery of over 100%.

MNE recovered 101.2% of the 1,800 nA·m CTF dipole, 26.3% of the 180 nA·m CTF dipole, and 72.3% of the 200 nA·m Elekta dipoles. Because P □ I², 26.3% amplitude recovery corresponds to 6.9% of the inferred power.

The lower-amplitude phantom conditions are closer to, but still substantially larger than, the per-vertex human estimates. These analyses therefore characterize possible amplitude bias at the tested phantom scales but do not directly calibrate the human resting-state estimates.

Across SNR settings, mean total current-dipole magnitude varied threefold, while recomputed total primary P varied 10.3-fold (Supplementary Figure 3).

Constraining dipoles normal to the cortical surface increased mean total current-dipole magnitude from 7.44 × 10□□ to 1.27 × 10□□ A·m and increased mean total primary P from 7.12 × 10□□ to 2.24 × 10□□ W. These effects remain within approximately one order of magnitude but are not negligible.

Together, these analyses show that the tested inverse-solution choices can shift the estimate by approximately one to two orders of magnitude; they do not directly calibrate or bound every source of bias.

To independently assess the order of magnitude of our resistance estimates, we implemented the dendritic-based resistance model described above. Across n = 30 subjects, we found the dendritic resistance was estimated at 185.2 Ω (range across individuals: 157.3–215.4 Ω), compared to a mean gray matter cortical-patch resistance of 914.1 Ω (range across individuals: 746.5–1089.7 Ω) (Figure 2D). Additionally, we observed limited spatial correspondences between the subject-averaged patch-derived versus neuron-count-derived resistances, after removing the shared area term in the denominator of both formulations (Spearman r=0.14).

**Figure 2:**
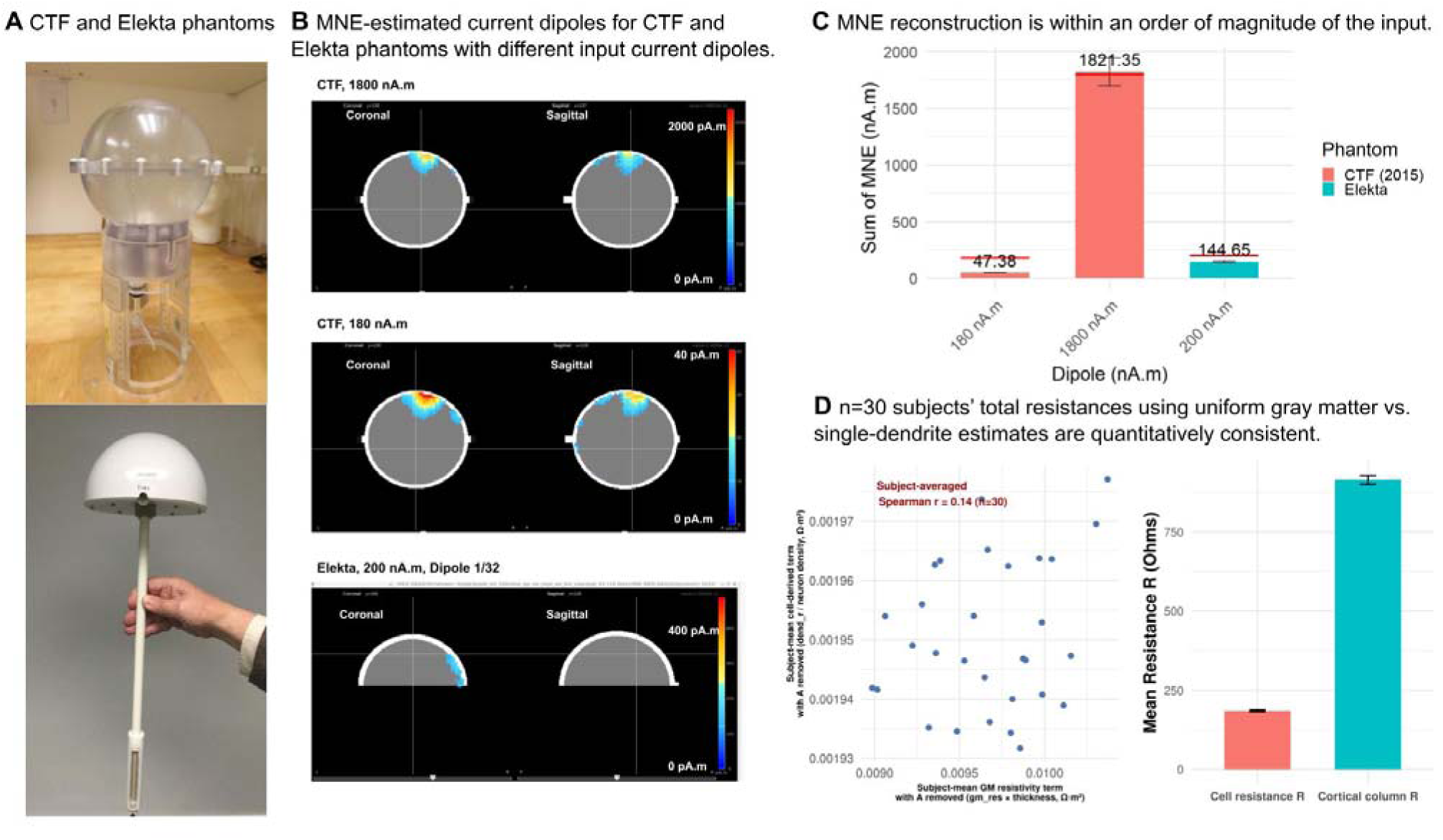
Order-of-magnitude checks of current-dipole and resistance estimates. Figure 2A shows the CTF (upper) and Elekta (lower) phantoms used to generate known current-dipole moments. Figure 2B shows representative MNE reconstructions, which were summed to produce the values in Figure 2C. Figure 2C shows the mean and standard deviation of total MNE recovery at peak amplitude (see Current-Dipole Recovery with Physical Phantoms). Figure 2D compares the gray-matter-resistivity resistance estimates with the neuron-count-based dendritic formulation across n=30 participants, including the spatial scatterplot and cohort means with standard errors.

These reformulations of the current-dipole and resistance estimates serve as order-of-magnitude checks within a simplified Ohmic circuit framework, which omits capacitive, frequency-dependent, and other dynamic tissue properties.

### Qualitative Cross-Model Comparison of Secondary-P Localization

To assess how the three-layer choice affected secondary-P localization, we compared a single simulated 20 nA·m somatosensory evoked field in the three-layer model for OMEGA participant sub-0002 with the same nominal source in a publicly available six-layer model from a different individual (https://zenodo.org/records/3888381; see Six-Layer FEM Model). Because the models differ in subject, anatomy, segmentation, and mesh construction, the comparison was treated descriptively.

In the six-layer single-source simulation, most of the secondary power was localized to gray matter. In the three-layer results, most secondary power was localized to the combined brain compartment, but the reported mean total secondary P was 1-2 orders of magnitude smaller than the total secondary P for the six-layer model. These descriptive values differ by roughly 27-fold; because they were obtained from different subjects and analysis inputs, their ratio is not a controlled quantitative model comparison.

Because the simulations used different subjects, imaging inputs, segmentation pipelines, mesh resolutions, and tissue definitions, this analysis does not provide a controlled quantitative validation of the three-layer model. It instead shows that both models localize most secondary dissipation to intracranial brain tissue, while leaving the magnitude and finer tissue distribution model-dependent.

For total secondary P, the three-layer model likewise localized most dissipation to the combined brain compartment (Supplementary Figure 2); this observation is descriptive and model-dependent.

### Secondary-Current Power Dissipation Under the RMS Approximation

Our initial calculations estimated power dissipation at cortical surface vertices on the FEM-derived model, but were limited to currents within gray matter directly beneath each vertex. To extend this approach, we used the full FEM model, representing the entire head volume as a conductive medium composed of discrete elements, each assigned a conductivity value based on tissue type (Figure 3C). This method allowed us to account for secondary currents dissipated by primary dipoles as they are driven throughout the head volume.

**Figure 3:**
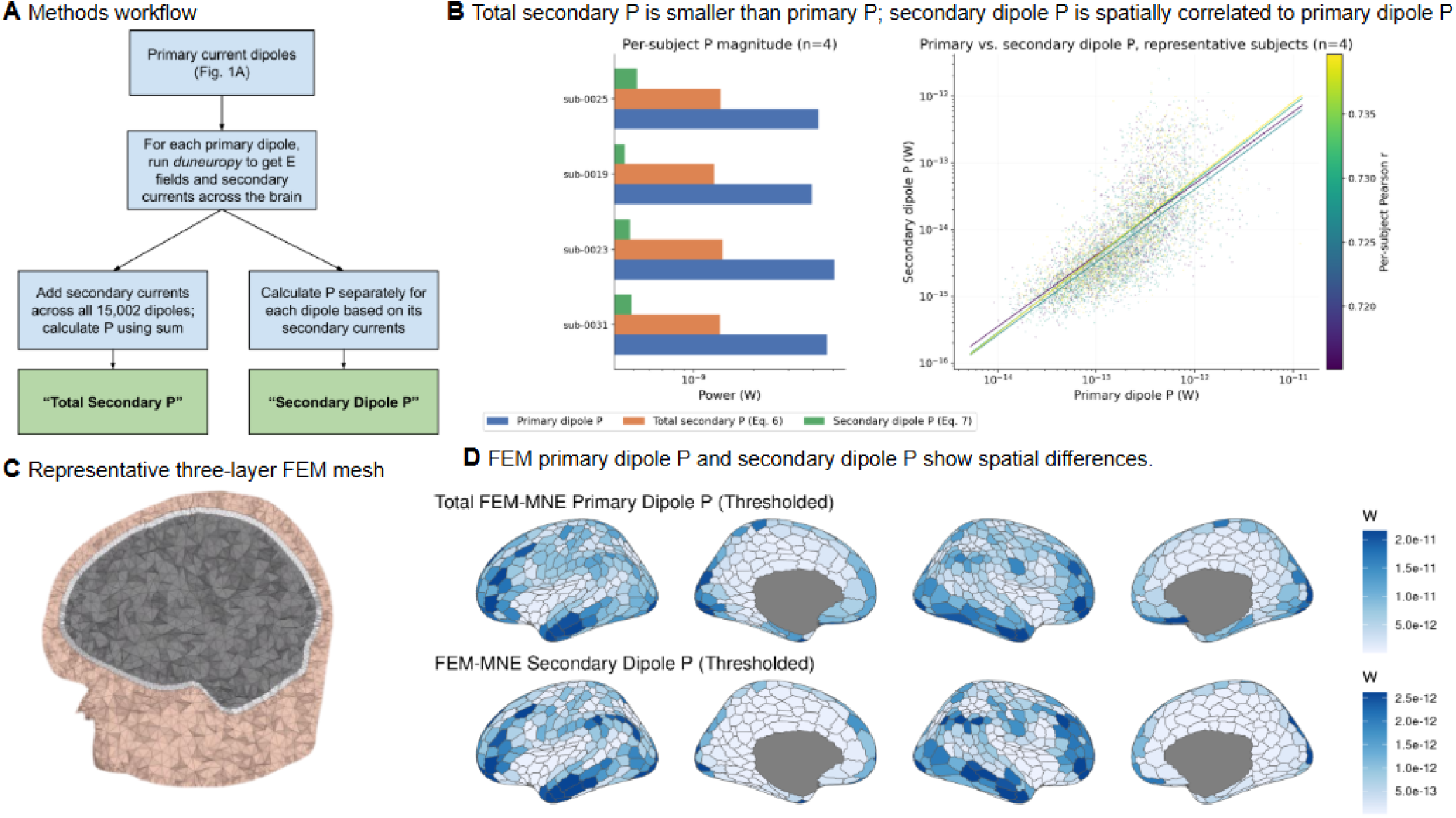
FEM derivations of secondary power consumption. Fig. 3A shows the methods workflow for deriving secondary dipole P and total secondary P, which are shown for each of our n=4 exemplar individuals in 3B. Fig. 3B shows the relative magnitude of primary dipole P, secondary dipole P, and total secondary P, as well as the correlation between primary and secondary dipole P across the n=4 exemplar subjects used in this analysis. Each point on the right-hand side of Figure 3B corresponds to a primary current dipole and a single cortical patch, and there is a separate fit shown for each participant (colored by correlation). Fig. 3C shows a representative FEM model used to derive the power estimations. Fig. 3D uses *ggseg* to visualize the spatial differences in the primary and secondary dipole P averaged across all individuals.

Secondary-current estimation is substantially more computationally expensive than primary dipole P, so we restricted this analysis to four successfully modeled exemplars (OMEGA IDs: sub-0019, sub-0023, sub-0025, and sub-0031; 2 women/2 men; all right-handed; mean age 26.7 years). The four exemplars had total current-dipole magnitudes close to those of the remaining cohort and highly similar parcellated spatial distributions. Their mean primary P was lower than that of the remaining participants but remained within the same order of magnitude. We therefore use them as central-tendency exemplars for the computationally intensive FEM analysis, not as a statistically representative subsample.

Under the RMS-source approximation, total secondary P was 27–32% of total primary P across these four exemplars (mean 30.3%). Thus, adding modeled secondary volume currents increased the estimate by approximately 30%, rather than by orders of magnitude. Spatial differences between primary and secondary dissipation are shown in Figure 3.

These FEM results indicate that modeled secondary currents under the RMS approximation do not bridge the gap with metabolic predictions; broader methodological and physiological differences remain.

### Power Correlations with Whole-Cortex and Within-Network Metabolism

After characterizing the magnitude and spatial specificity of the MEG-derived estimates, we investigated their relationship with cortical metabolism. Primary P maps were parcellated into the Schaefer-600 atlas and compared with PET-derived glucose and oxygen metabolism maps (Vaishnavi et al., 2010; Markello et al., 2022) and with transcriptomic maps of 34 metabolic processes derived from the Allen Human Brain Atlas (Pourmajidian et al., 2025). We repeated all spatial correlations using the MNE current-dipole maps for descriptive comparison.

MEG-derived power correlated with oxygen metabolism (ρ = −0.147, p = 0.0047); the current dipole also correlated with oxygen metabolism (ρ = −0.132, p = 0.0110). The power coefficient was numerically larger in magnitude, although we did not formally test the difference between the two dependent correlations. Neither metric correlated significantly with glucose metabolism (power: ρ = 0.039, p = 0.5493; current dipole: ρ = 0.043, p = 0.5109; Figure 4A).

**Figure 4:**
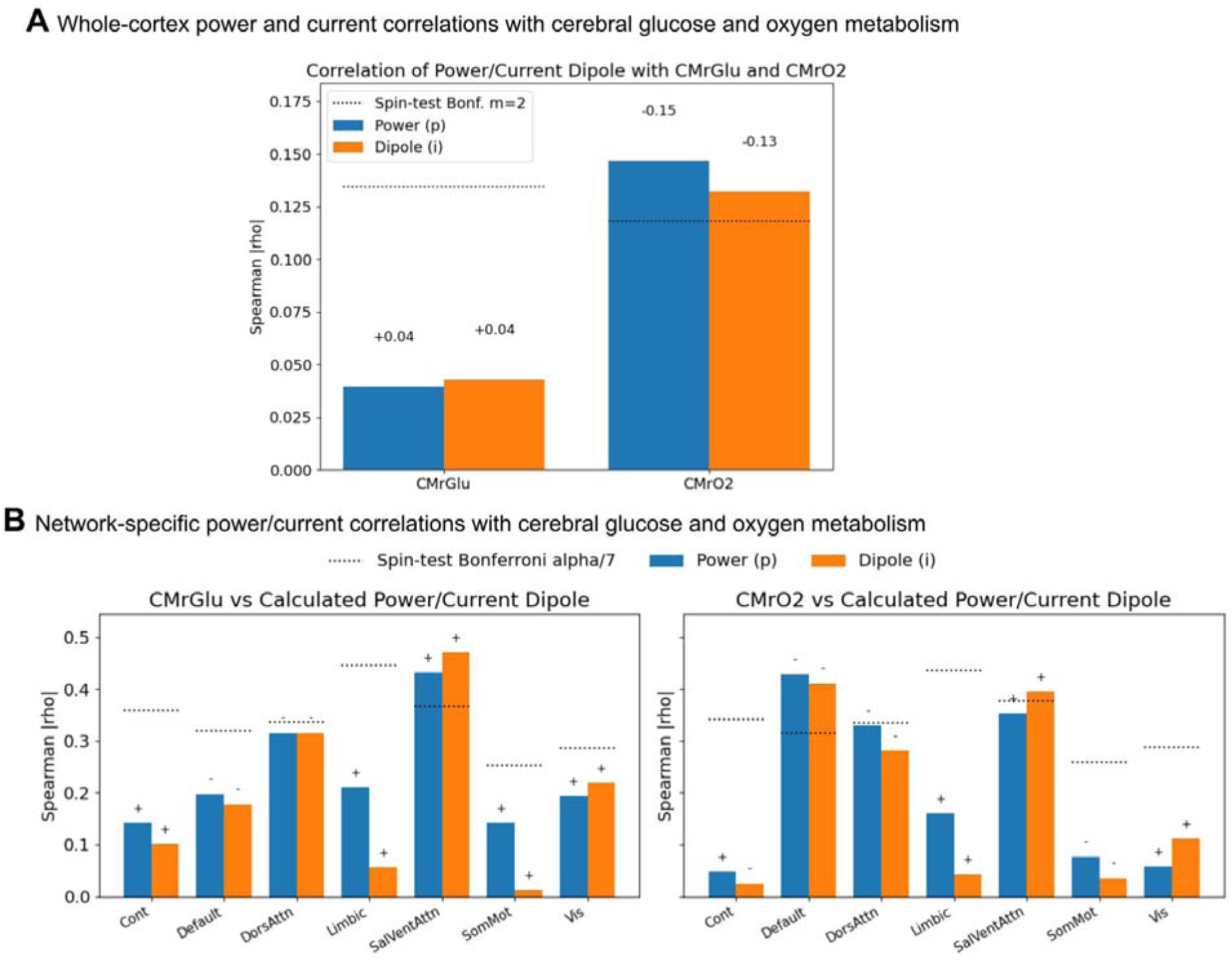
Correlations between parcellated power dissipation and PET-derived metabolic maps across the cortex and within functional networks; signs above bars indicate correlation direction. Figure 4A compares primary P and current-dipole maps with glucose and oxygen metabolism across Schaefer parcels. Figure 4B shows the corresponding within-network correlations defined by Yeo et al. (2011). Each network has a different critical Spearman coefficient because parcel counts and spatial-null distributions differ. Abbreviations: CMrGlu, cerebral metabolic rate of glucose; CMrO, cerebral metabolic rate of oxygen.

In the ventral attention network, both MEG-derived power (ρ = 0.432, p = 0.0010) and the current dipole (ρ = 0.470, p = 0.0002) correlated positively with glucose metabolism and exceeded the Bonferroni-corrected threshold (α = 0.05/7 = 0.00714; Figure 4B).

For oxygen metabolism, power correlated significantly in the default-mode network (ρ = −0.428, p = 0.0002) and dorsal-attention network (ρ = −0.329, p = 0.0071). In the default-mode network, the current-dipole coefficient was ρ = −0.411 (p = 0.0002). In the ventral-attention network, the power coefficient (ρ = 0.353, p = 0.0140) did not reach Bonferroni significance, whereas the current-dipole coefficient did (ρ = 0.395, p = 0.0034). The absolute correlation coefficient for power was numerically larger than that for the current dipole in 9 of 14 network-by-metabolite comparisons. This is a descriptive comparison and should not be interpreted as evidence of a significant difference between metrics.

Ketone-body utilization and ketone metabolism showed the largest absolute whole-cortex correlations with power (ρ = 0.270 and 0.256), compared with ρ = 0.213 and 0.210 for the current dipole. Power also showed numerically larger absolute correlations in 6 of 7 and 5 of 7 networks, respectively (Figure 5; Supplementary Table 1).

**Figure 5:**
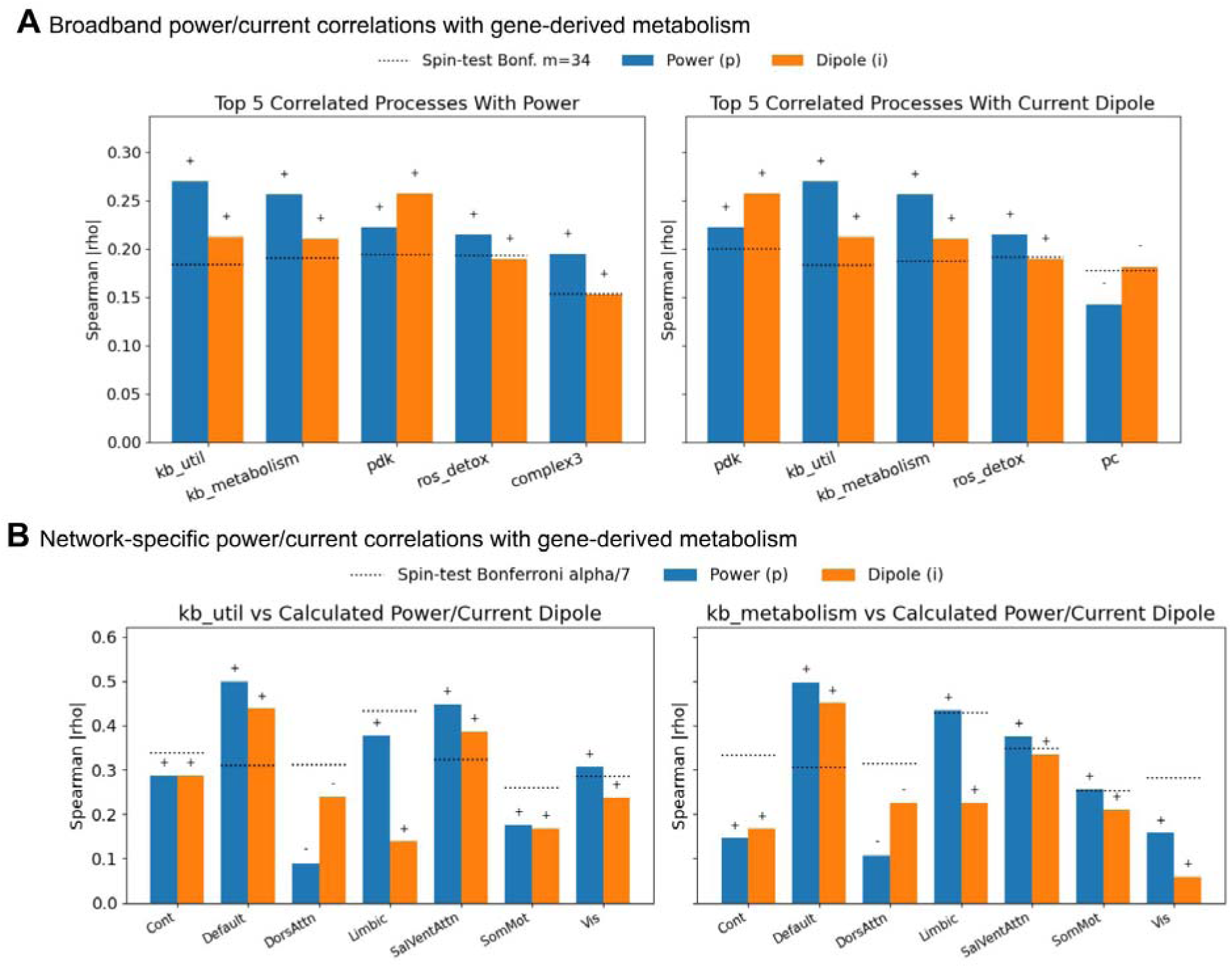
Correlations between parcellated power dissipation and gene-set-derived metabolic maps across the cortex and within functional networks. (Pourmajidian et al., 2025), displayed as in Figure 4. Figure 5A compares primary P and current-dipole maps with the top five of 34 gene-set-derived metabolic processes. Figure 5B shows associations with ketone-body utilization and ketone metabolism within Yeo networks. Abbreviations: kb_util, ketone-body utilization; kb_metabolism, ketone-body metabolism; pdk, pyruvate dehydrogenase kinase; complex3, Complex III of the electron transport chain; ros_detox, detoxification of reactive oxygen species; pc, pyruvate carboxylase; mas, malate–aspartate shuttle; atpase, sodium/potassium ATPase pump.

The associations remained significant after controlling separately for cortical thickness, support area, and the combined geometric factor 1/(LA). The oxygen-metabolism coefficient became more negative after controlling for geometry, suggesting that the association is not explained by L or A alone. This result does not, however, establish that the remaining association is specifically driven by the squared current term (Supplementary Figure 5).

Taken together, the analyses showed modest, spatially heterogeneous correspondences between MEG-derived power and selected metabolic maps. Numerical differences between power and current-dipole coefficients were descriptive because the dependent correlations were not formally compared.

## Discussion

### Discrepancy Between MEG-Derived and Metabolic Power Estimates

We sought to estimate the electrical power dissipated by synchronized cortical activity detectable with resting-state MEG. Across the primary-analysis cohort, mean total primary P summed over the modeled cortical source space was 6.6 × 10 W, several orders of magnitude below metabolic power estimates and model-based estimates of cortical computation. This value should not be interpreted as total cortical power: MEG is insensitive to asynchronous, geometrically opposed, and otherwise sensor-invisible source configurations.

Targeted phantom, inverse-solution, resistance, and FEM analyses indicated that the methodological choices tested here can alter the estimate by approximately one to two orders of magnitude. These effects are important but insufficient to bridge the full gap with metabolic benchmarks. We therefore interpret the result as an empirical lower-bound estimate of power associated with MEG-visible synchronized activity.

We acknowledge that, due to signal cancellation, MEG is inherently limited in capturing the full magnitude of cortical sources. MEG is sensitive to pyramidal neurons that are spatially coherent and similarly aligned relative to the cortex (an “open field”), but incapable of measuring neurons whose geometric orientations are misaligned to each other (a “closed field”), since their dipoles cancel at the level of a cortical patch (Lopes da Silva et al., 2010). Hence, we recognize that, due to this signal cancellation, our approach cannot calculate the total amount of power emitted by cortical neurons, outside of very specific scenarios where all pyramidal cells are synchronized (e.g. an epileptic seizure). In addition to signal cancellation due to a lack of synchrony, there is also signal cancellation due to the spatial opposition of MEG sources, such as sources directly facing each other on sulcal walls. Finally, the macroscopic detectability of MEG sources depends on the sources’ relative orientation to the head and sensors; radial gyral sources are often harder to detect than sulcal sources.

For these reasons, we interpret the metric as a lower bound on power dissipated by MEG-visible synchronized cortical currents. When parcellated into brain regions, the metric showed statistically significant but modest spatial associations with PET- and gene-derived measures of cortical metabolism.

### Methodological Considerations

Our analysis uses a purely resistive Ohmic approximation. Under the quasi-static approximation conventionally used in MEG, inductive effects are neglected; however, membrane capacitance, frequency-dependent impedance, cellular heterogeneity, and recurrent connectivity are not represented. Each vertex-associated cortical patch is also treated as an independent effective circuit. These assumptions limit the absolute biophysical interpretation of the estimate.

The primary analyses used SNR = 3 and unconstrained dipole orientation. Sensitivity analyses showed that varying SNR from 1 to 5 changed total primary P by approximately one order of magnitude, while constraining dipoles normal to the cortical surface changed total primary P by approximately threefold. These effects demonstrate that inverse-solution assumptions materially affect amplitude, although they do not account for the several-orders-of-magnitude difference from metabolic benchmarks.

The three-layer FEM model combines gray matter, white matter, and CSF into a single brain compartment. A separate six-layer, single-source simulation suggested limited CSF dissipation for that particular source, but the comparison used a different subject, anatomy, segmentation, and mesh. It should therefore be interpreted qualitatively and cannot establish the CSF contribution across source locations.

### Physiological Distinctions Between Electrophysiological and Metabolic Measures

Beyond modeling assumptions, fundamental physiological differences may explain why MEG-derived power dissipation diverges from metabolic estimates. MEG primarily detects large-scale, synchronous cortical currents in superficial layers (Baillet, 2017; Baillet, Mosher, & Leahy, 2001). In contrast, the majority of metabolic consumption arises from processes invisible to MEG, including synaptic vesicle cycling, neurotransmitter reuptake, and the maintenance of resting membrane potentials (Alle et al., 2009; Castrillón et al., 2023). Additionally, MEG captures neural dynamics at the millisecond timescale, whereas PET provides time-averaged measures of biochemical energy use over tens of minutes (Clarke & Sokoloff, 1999; Vaishnavi et al., 2010). These differences in spatial and temporal sensitivity contribute to the observed mismatch in magnitude.

### Regional Correspondences With PET and Gene Expression

After spatial-null correction, MEG-derived power showed modest associations with whole-cortex oxygen metabolism, glucose metabolism in the ventral attention network, and oxygen metabolism in the default-mode and dorsal-attention networks. The corresponding power and current-dipole coefficients were generally close in magnitude, and their differences were not formally tested.

Ketone-body utilization and ketone metabolism were the strongest transcriptomic correlates of the power map at the whole-cortex level. Power also showed numerically larger coefficients than the current dipole in most networks, including the default-mode network.

These exploratory spatial associations do not demonstrate preferential ketone utilization or a shift in metabolic substrate. They require validation using electrophysiological and metabolic measurements acquired in the same individuals.

### Implications and Future Directions

Taken together, our results provide an empirical lower-bound estimate of electrical power dissipation associated with MEG-visible synchronized cortical activity. Although incomplete in absolute terms, the regional metric may prove useful for comparing conditions characterized by altered cortical synchrony, provided that its reliability, sensitivity, and metabolic interpretation are established in independent cohorts.

Future work should refine the electrical model by incorporating capacitive and frequency-dependent tissue properties; combine electrophysiological and metabolic measurements within the same participants; and evaluate the reliability and sensitivity of the regional metric across lifespan, disease, and intervention.

By relating a physically defined electrophysiological power metric to regional metabolism, this study provides a first step toward multiscale models linking synchronized cortical activity with biochemical energy use.

## Acknowledgements

The Elekta phantom data was provided by Ken Taylor and John Mosher (Epilepsy Center, Cleveland Clinic Neurological Institute, Cleveland, OH USA). The CTF phantom data was provided by Elizabeth Bock, Francois Tadel and Sylvain Baillet (MEG Lab, McConnell Brain Imaging Center, Montreal Neurological Institute, McGill University, Canada). We would like to thank Moohebat Pourmajidian for her input on extracting and processing the gene-set derived metabolism maps.

## Code availability

Code for all analyses is available at https://github.com/vik16nathan/meg_power.

