## Supplementary information for "Electrophysiological “Brain Power” Dissipation: MEG Lower-Bound Estimates and Regional Metabolic Correlates"

**Supplementary Materials**

**Supplementary Methods 1:** Temporal Aggregation is Equivalent to using Root-Mean-Square Current Dipole for Primary Dipole P

To calculate the average primary dipole P for a single vertex across all T time-points:

$P_{avg} =\frac{1}{T}\sum_{t=1}^{T} \rho{D^{2}/(L\cdot A)}$

Rearranging, and substituting the magnitude of current dipole D as $D = \sqrt{(D_{x}^{2}+D_{y}^{2}+D_{z}^{2})}$

$P_{avg}=\sum_{t=1}^{T} {{\sqrt{\frac{1}{T}(D_{x}^{2}+D_{y}^{2}+D_{z}^{2})}}^{2}\rho/(L\cdot A)}$

Assuming that ⍴, L, and A are time-invariant, and further rearranging:

$P_{avg}= {{\sqrt{\frac{1}{T}\sum_{t=1}^{T} (D_{x}^{2}+D_{y}^{2}+D_{z}^{2})}}^{2}\rho/(L\cdot A)}$

Hence, applying the definition in Eq. (8) from the main text, where || || represents the L2 norm,

$P_{avg}= {{{|\left| D \right||}_{rms, v}^{2}}\rho/(L\cdot A)}$

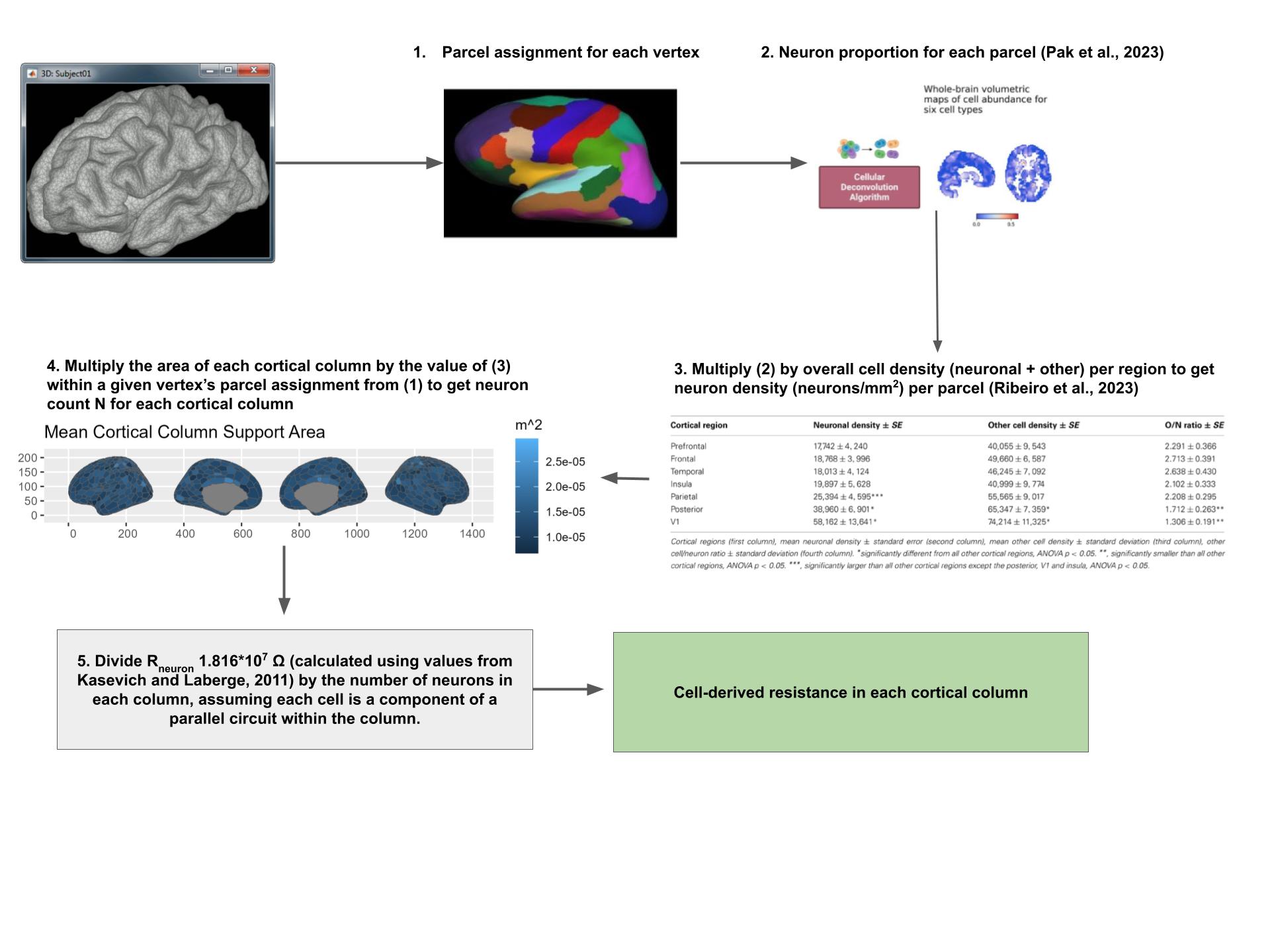

**Supplementary Figure 1:** Schematic showing the steps required to calculate the neuronal count-derived resistance underlying each vertex/cortical column.

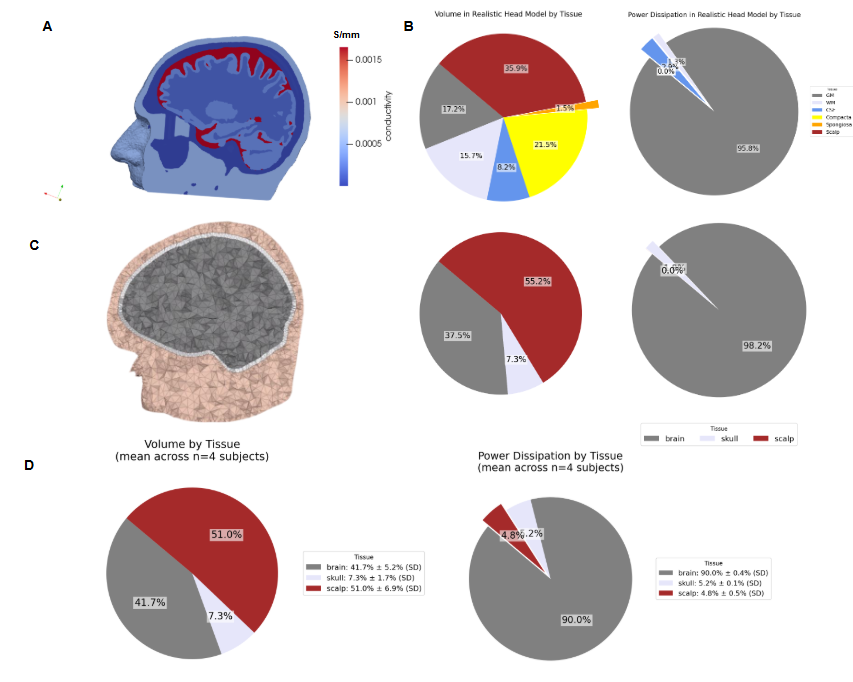

**Supplementary Figure 2:** Secondary power dissipation across tissue types in a more realistic six-layer FEM model is qualitatively similar to the three-layer FEM model used in our analyses. The realistic FEM mesh is shown in 2A and the volume and power outputs across tissue types in each FEM mesh are shown in 2B; the total P was 7.785 × 10^-8^ W. This analysis was conducted by simulating the currents induced by a single somatosensory evoked potential (SEP) of 20 nA.m located on the postcentral gyrus. Fig. 2C indicates that most of the secondary power dissipation from the same 20 nA.m SEP is localized to the brain, despite most of the volume corresponding to scalp tissues; the total secondary P is 2.935×10⁻⁹ W. Figure 2D extends this finding to the resting-state brain activity, tracking the total secondary P induced by the RMS current dipoles over 120 s of resting state activity within each FEM element, shown as an average across n=4 representative subjects. In the three-layer results, most secondary power was localized to the combined brain compartment, and the reported mean total secondary P across four participants was 1.73 × 10⁻⁹ W.

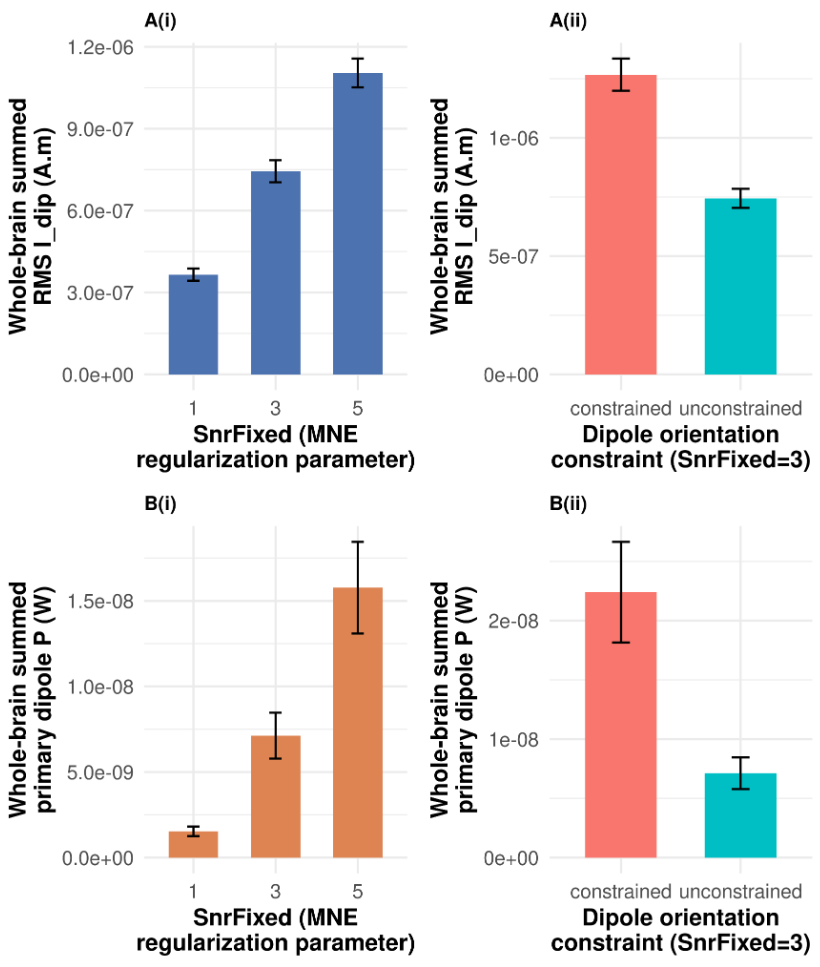

**Supplementary Figure 3:** Sensitivity analyses show similar order of magnitudes for our minimum-norm estimation (MNE) (A), when varying: (i) the signal-to-noise ratio (SNR) regularization parameter and (A) (ii) whether the sources were constrained to be normal to the cortical surface, or unconstrained to estimate the x, y, and z components of each current dipole. In (A) (i), as expected, we see that a larger SNR parameter results in larger estimates of primary current dipoles, whereas in (A) (ii), we see that constraining the currents normal to the cortical surface results in a larger current dipole estimate. In (B), we apply these currents to the primary dipole P, showing a greater divergence in P when calculating it across (i) different SNR parameters (ii) constrained vs. unconstrained orientation, but the same qualitative trends as the current dipole.

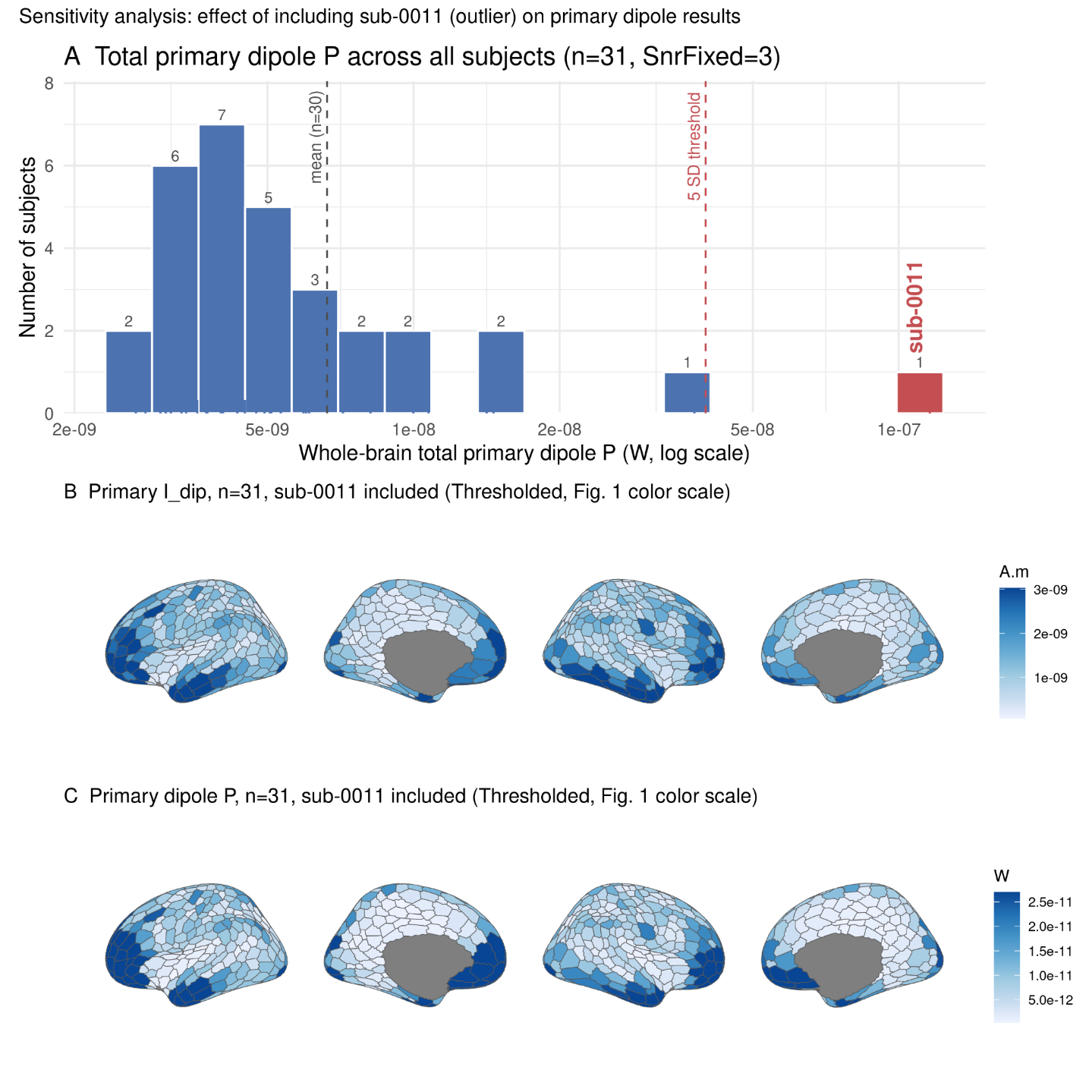

**Supplementary Figure 4**: We confirm that sub-0011 is an outlier using (A) a histogram of log-scaled total primary dipole P, with the 5 SD outlier threshold labelled in red. In (B) and (C), we show the average primary current dipole across all n=31 subjects, but using the same colorbar scale as the n=30 subjects in Fig. 1 to enable direct comparison between maps.

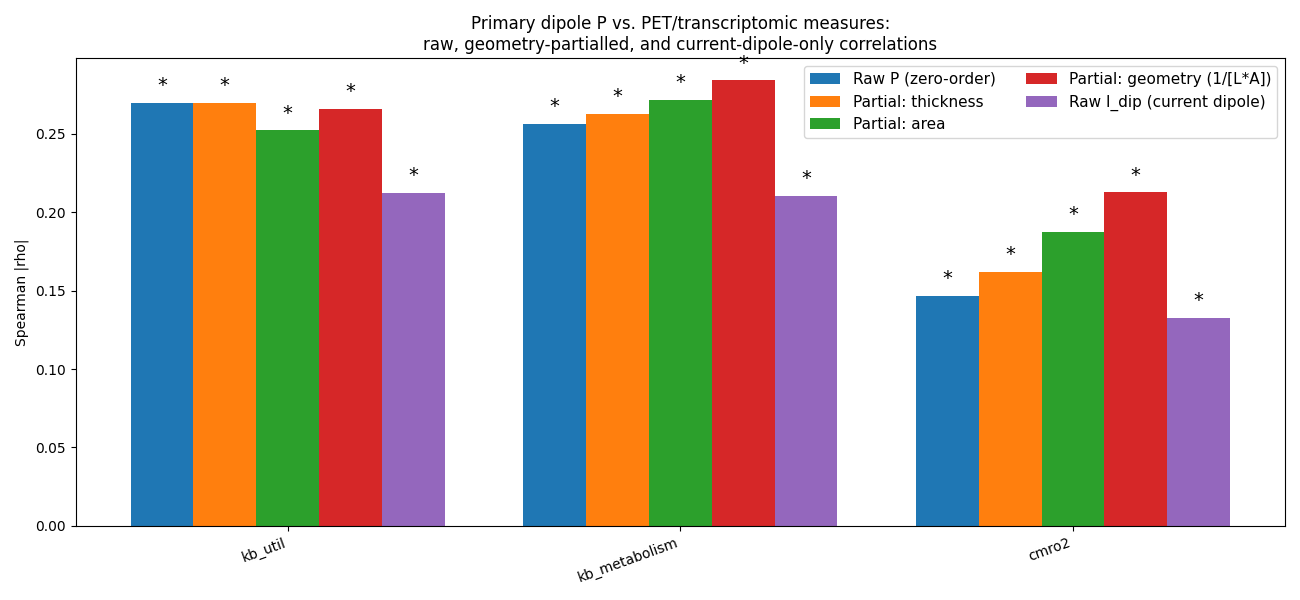

**Supplementary Figure 5:** We show that, after partial correlations correcting for cortical thickness (L), vertex support area (A), and their combination in our primary dipole P formula (1/(L × A)), that our primary dipole P metric still shows slightly higher correlations with the “top” metabolism maps than the primary current dipole. We show the results of the partial correlation analysis for maps of  ketone body utilization, metabolism, and CMrO_2_, using * to represent significance under the same Bonferroni-corrected, spatial-null model-derived correlation threshold used for these maps in Figures 4 and 5.

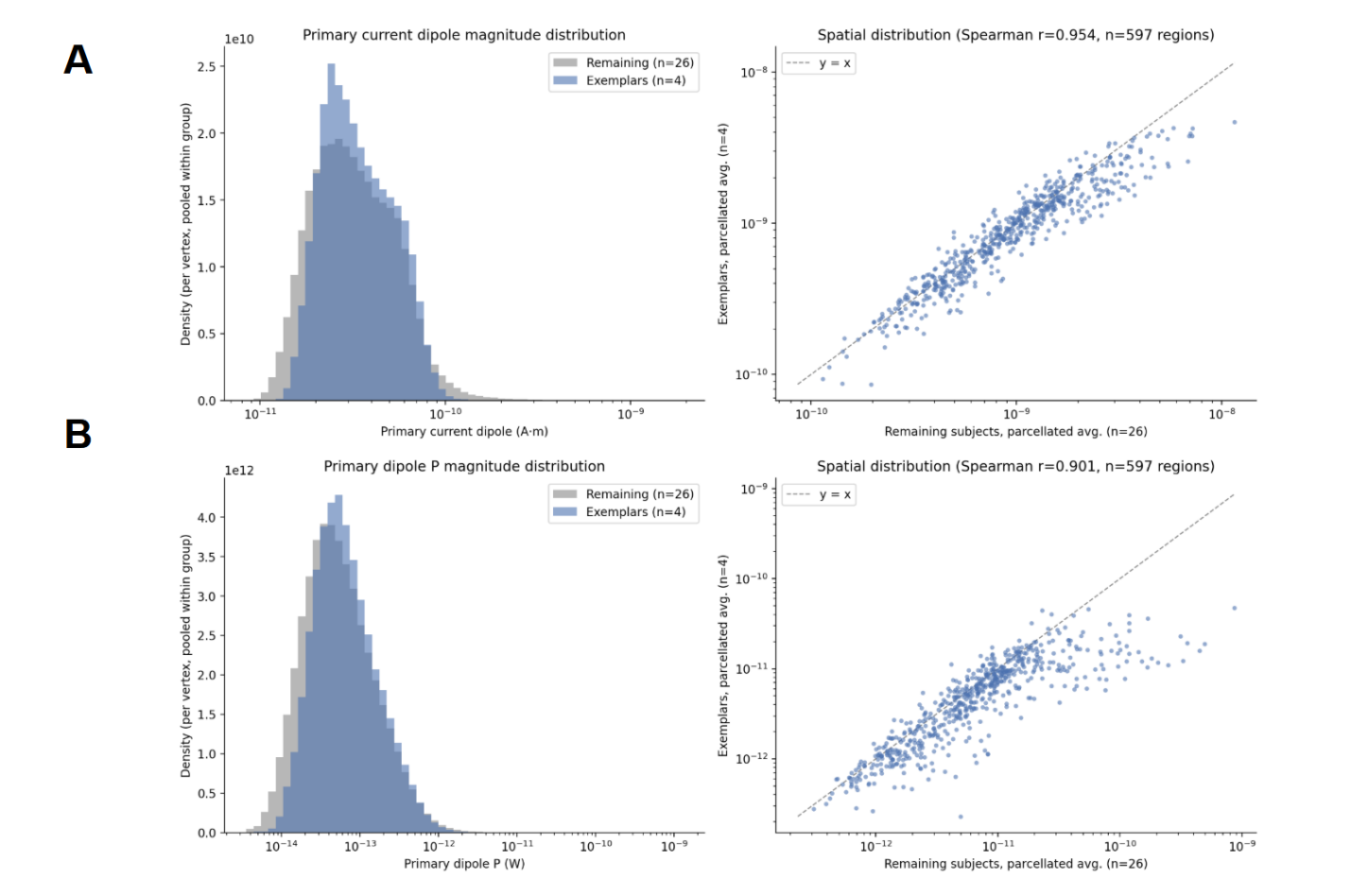

**Supplementary Figure 6:** We compare the distributions between our four central-tendency exemplar subjects (sub-0019, sub-0023, sub-0025, sub-0031) versus to the rest of the OMEGA cohort used in this paper.

**Supplementary Information: Idealistic Single-Neuron Power Consumption Scaled Across the Brain**

After validating the physical approaches used to calculate our initial lower bound, we sought to compare our estimates against a bottom-up calculation of whole-cortex power; namely, a calculation that used Ohm’s law with the current dipole and resistance of a single neuron to calculate power, before multiplying this power by the number of neurons in the cortex. In place of an empirical dipole, we used the estimated dipole moment of the envelope of action potentials from a single layer V pyramidal cell, which was simulated to be 0.29 pA.m (Murakami and Okada, 2006). We use this as the value of I_dip_ for a single neuron. We then calculated the cellular power per neuron using P=( I_dip_ /L)^2^ R_neuron_ , with L= 1,200 µm and R_neuron_ =1.816*10^7^ Ω previously calculated from the values in Kasevich and Laberge, 2011 (*Methods*). Finally, we multiplied this power by the total number of dendrites in the human cortex, which we assume is upper-bounded by the number of neurons in the cerebral cortex (16 billion; Herculano-Houzel, 2009). This calculation resulted in a whole-cortex power estimate of 0.0170 W.

We find our cortical patch-derived estimates to be far smaller than this bottom-up estimate, underscoring the divergence in the theoretical vs. empirically estimated amount of power that can be attributed to the MEG signal. This bottom-up model assumes that all cortical neurons emit this strong dipole, driven by sodium ion influx at a given moment in time; in fact, it is widely argued that MEG cannot measure transient, sodium-spike-driven brain activity due to the variability in timing of these events (“temporal jitter”) (De Munck et al., 1992). We also note that Murakami’s estimate is based on a theoretical model of a compartmentalized neuron, and thus cannot be used as an empirical validation. Additionally, Murakami’s estimate for current dipoles is 20-45 times larger than an earlier estimate using the classical, cylindrical model of a neuron (Hämäläinen et al., 1993). We instead view this estimate as a theoretical upper bound for the power consumption that can be tracked by a MEG scan at a given moment in time.

**Supplementary Table 1:** Whole-brain correlations between MEG-derived current and power, and all the metabolic processes in Supplementary Figure S9 of Pourmajidian et al., 2025. See Pourmajidian et al., 2025 for full names of the abbreviated metabolic processes shown below.

| **Pathway** | **Power correlation** | **Current dipole correlation** |
| --- | --- | --- |
| kb_util | 0.2697 | 0.2125 |
| kb_metabolism | 0.2564 | 0.2104 |
| pdk | 0.2223 | 0.2574 |
| ros_detox | 0.2150 | 0.1901 |
| complex3 | 0.1953 | 0.1528 |
| complex1 | 0.1908 | 0.1375 |
| etc | 0.1742 | 0.1134 |
| oxphos | 0.1723 | 0.1150 |
| atpsynth | 0.1713 | 0.1358 |
| kb_synth | 0.1678 | 0.1472 |
| pc | -0.1424 | -0.1815 |
| pdp | 0.1401 | 0.1432 |
| gln_glu_cycle | -0.1326 | -0.1488 |
| betaox | 0.1223 | 0.1461 |
| lactate_transport | -0.1075 | -0.1239 |
| glycolysis | 0.1021 | 0.0539 |
| complex2 | 0.0921 | -0.0074 |
| mas | -0.0900 | -0.1312 |
| pdc | 0.0770 | 0.0745 |
| complex4 | 0.0756 | 0.0150 |
| gps | -0.0635 | 0.0139 |
| tca | 0.0622 | -0.0198 |
| fa_metabolism | 0.0561 | 0.0607 |
| fa_synth | -0.0503 | -0.0715 |
| glycogen_synth | 0.0444 | 0.0083 |
| atpase | -0.0440 | -0.1105 |
| ros_gen | 0.0386 | 0.0264 |
| ppp | 0.0385 | -0.0476 |
| lactate_metabolism | 0.0239 | -0.0291 |
| creatine | -0.0189 | -0.0818 |
| no_signalling | -0.0133 | 0.0392 |
| glycogen_metabolism | 0.0046 | -0.0429 |
| glycogen_cat | -0.0028 | 0.0353 |
| lactate | 0.0020 | -0.0480 |
